# Detecting ecosystem trends in response to climate and disturbance across continental plot networks: a power analysis

**DOI:** 10.1101/2024.10.14.618336

**Authors:** Greg R. Guerin, Irene Martín-Forés, Rachael V. Gallagher, Samantha E.M. Munroe, Ben Sparrow

## Abstract

Plant communities are dynamic: cyclical, directional and stochastic changes in species composition make interpretation of responses complex. The multi-dimensionality of ecological change is reduced by ecosystem indicators that quantify functional responses to environmental drivers. However, thresholds of detectable change in indicators may vary by ecosystem or the spatial configuration of monitoring sites within regions.

We quantified the power of a continental terrestrial ecosystem plot network (TERN Ausplots, Australia) to detect change in ecological indicators linked to disturbance and climate: community temperature index (CTI), proportional abundance of photosynthetic pathways (C_4_%), and species abundance distribution (SAD). We simulated trend and noise scenarios and calculated power and effect size across spatial clusters of plots using linear mixed effects models. We then assessed factors that influenced change detection capacity.

Power varied substantially across the network. Trends with a minimum magnitude of 0.5–3.0°C in CTI, 0.01–0.3 in C_4_% and 0.1–0.5 in the SAD sigma were detectable, depending on cluster identity and noise. Increased plot replication within a spatial cluster increased power and lower variance among baseline replicates increased effect size for CTI and C_4_% but not SAD. Latitudinal patterns in change detection capacity emerged for CTI and C_4_%. The effect size of increasing CTI was greater in the tropics due to lower among-plot variance, while power to detect increasing C_4_% was higher in the temperate south, where C_4_ is rare.

Indicators reveal trends in complex ecological data that can inform decision-making. Increasing spatial replication and minimising spatial variation by stratifying plots to the same ecological community enhance change detection, albeit with different effectiveness by indicator. The results suggest change detection differs by location, irrespective of sampling factors such as number of plots. Strategies and thresholds for change detection may therefore apply unequally across environments and should be considered by region and response parameter.

## Introduction

Terrestrial plant communities exist in a state of flux (Link *et al*. 2010). Species composition, vegetation cover, structure and primary productivity, for example, are influenced by climatic fluctuations, historical and biogeographical factors (e.g., migration barriers, imprints of past climate), ecological drift, and disturbance from land-use changes, grazing, cyclones, floods or fire (Philippi *et al*. 1998; Vesk *et al*. 2004; Chase 2007; Ibanez *et al*. 2019; Keith *et al*. 2022). Climate change and altered disturbance regimes are key threats to ecosystems that operate against this background of standing variability (Greenville *et al*. 2012; Kampichler *et al*. 2012).

The response of biota to these disturbance drivers can be detected through monitoring, which is vital for informing land management, determining conservation priorities, and carbon accounting (Nolan *et al*. 2018; Baker *et al*. 2019; Oliva *et al*. 2020; Sparrow *et al*. 2020a; Baruch *et al*. 2022). However, raw observations of ecosystem status and trends can be complex or difficult to interpret, particularly multivariate changes to species composition and relative abundance (Doren *et al*. 2009). Across a monitoring plot network, numerous species that respond individually to multiple simultaneous drivers may be observed (Philippi *et al*. 1998). Indeed, individual species-based approaches can fail to predict higher-order responses to environmental change (Vesk *et al*. 2004). Change detection across a complex system of species is further hampered by practical constraints such as imperfect species detection (Clarke *et al*. 2012; Baker *et al*. 2019) and logistic constraints to the timing of revisits across a national network of sites (Martín-Forés *et al*. 2021).

Novel suites of monitoring networks are attempting to address and synthesise knowledge gaps about ecological change at increasing large scales (Schimel 2011). Often, the initial focus of national and continental status and trends monitoring networks is on the immense task of ensuring environmental coverage and representation (Hoffman *et al*. 2013; Metzger *et al*. 2013; Guerin *et al*. 2020). The power of these networks to detect change has rarely been quantified despite being an equally important role of a successful observation network (Buckland & Johnston 2017; O’Hare *et al*. 2020; Guerin et al. 2021a). Two challenges arise for plot networks maturing from an establishment phase into an operational monitoring system. The first is to switch from a focus on environmental coverage and survey methods to a focus on quantifying thresholds for detectable change. Many networks have unknown power to detect change regionally (Buckland & Johnston 2017; Baker *et al*. 2019), due to a focus on initial design (O’Hare *et al*. 2020). TERN Ausplots in Australia (Sparrow et al. 2020b) is no different. The power analysis reported here seeks to retrospectively assess change detection capacity, a valuable review process that can inform network redesign and monitoring protocols going forward (O’Hare *et al*. 2020). The second challenge follows from the first and lies in harnessing complex observations to interpret trends (Baruch *et al*. 2022). For TERN Ausplots, this challenge is exemplified by high species turnover (Baruch *et al*. 2018) with many individual trends at plot level (Baruch *et al*. 2022), combined with an absence of formal replication of plots targeting particular ecological communities, and no fixed schedule of revisits, leading to a patchwork of ecological community samples forming units of change detection.

One approach to reducing the complexity of species occurrence information in a monitoring framework is to use ordination techniques to identify major gradients in composition, which can then be related to particular environmental parameters (Philippi *et al*. 1998). In this way, trajectory through composition space can be tracked over time and associated with variables such as rainfall or time since fire (Clarke *et al*. 2005). An alternative dimension reduction approach to ordination is to calculate univariate indices that serve as indicators of underlying species composition and that can be correlated to key environmental parameters or changes of interest. Such ecosystem indicators can be calculated from complex underlying observations of the structure or composition of ecological communities. Indicators synthesise raw data with the potential to link ecological trends to specific environmental changes (Matthews & Whittaker 2015), bridging theory, monitoring and management (Link *et al*. 2010). Tractable indicators are suitable for environmental reporting and improve early warning and interpretation of changes detected through systematic monitoring (Doren *et al*. 2009). Understanding how aggregated community level traits respond to local disturbance and environmental drivers is critical for detecting and disentangling responses to climate change (Bruelheide *et al*. 2018). Useful indicators, however, must be measurable and sensitive to change (Link *et al*. 2010). Indicators ideally have specific drivers (e.g., temperature, grazing regime or recovery from logging, storms, fires and drought; Link *et al*. 2010), but these relationships are often poorly understood, particularly in Australian drylands (DeMalach *et al*. 2019).

The capacity to detect trends in suitable indicators can be quantified before applying them to the interpretation of on-ground monitoring data. Two statistical aspects of change detection are power (probability of detecting a given trend) and effect size, defined here as proportion of variance explained by that trend. Ideally, a monitoring network has high power to detect ecologically significant changes early enough to highlight and respond to those changes before they become extreme (Bellingham *et al*. 2020). Ability to detect a particular trend is also affected by the level of existing variation within the ecosystem and the ratio of noise to signal in the temporal data (i.e., correlation of trends among sites) due to competing drivers, random changes and sampling error (Philippi *et al*. 1998; Barnett *et al*. 2019; Oliva *et al*. 2020). The magnitude of a trend relative to standing variation and noise therefore influences certainty that the detected trend is both real and ecologically important, given the milieu of spatial and temporal ecological dynamics. Standing variation can be at least partially controlled through sampling design that stratifies spatially replicated observations to similar systems. The magnitude of temporal noise (i.e., non-trend fluctuation at the local scale) is expected to be highly variable through time and space and can be minimised through rigorous and standardised monitoring protocols to reduce sampling error (Sparrow *et al*. 2020b).

We investigated which sampling regimes increase change detection power and explored minimum detectable levels of change in species composition relative to environmental drivers. To do so, we tested continental-scale sensitivity of ecosystem indicators to directional change, making use of new national datasets that include standardised measurements of plant species abundance, and matching trait information. Using data from >700 surveys of monitoring sites across Australia from TERN Ausplots (Sparrow et al. 2020b), we enrich species abundance data with functional trait information to calculate ecological indices reflecting, respectively, species dominance, thermal tolerances, and photosynthetic capacity, which are critical attributes underlying vegetation function (Harrison *et al*. 2010). Observed and simulated changes in these indicators across the Ausplots network are used as the basis for understanding sampling—power relationships.

Questions:

- What is the effect size of simulated directional change in three indicators across a plot network (i.e., how much variance is explained by the trend): community temperature index (CTI), proportional abundance of C_4_ species (C_4_%), and species abundance distribution (SAD)?
- How do power and effect size vary according to the number, spatial location, and variance among spatially replicated baseline plots?
- What kind of additional spatial replication will provide greatest gains in change detection across plot networks?
- What thresholds of change are detectable given non-directional ‘noise’ in species composition?

## Methods

### Species relative abundance dataset

Species abundances were scored across Ausplots, the TERN Ecosystem Surveillance network of monitoring plots, which sample the terrestrial habitats of Australia (Fig. 1). TERN Ausplots is a nationally distributed network of one-hectare vegetation monitoring plots representing Australia’s main vegetation systems including eucalypt and acacia woodlands and shrublands, tussock and hummock grasslands and chenopod shrublands (Guerin *et al*. 2017; Guerin *et al*. 2020; Sparrow *et al*. 2020b). The sites span the full latitudinal range of the continent from the island of Tasmania in the south to Cape Yorke Peninsula in the north and cover the diversity of terrestrial environments. The plot network has been stratified and gap-filled to be representative of major environments and vegetation types and also incorporates several climatic transects (Guerin *et al*. 2019; Guerin *et al*. 2020; Guerin *et al*. 2021a). The initial focus on capture of environmental and ecological variation has meant that the network lacks a formal framework for statistical change detection such as might include replication for statistical validity and schema for periodic revisit panels that are unbiased and representative (van Dam-Bates *et al*. 2018; O’Hare *et al*. 2020). At each plot, the cover of vascular plant species is measured using 1,010 point-intercepts arranged along a standardised grid of 10 transects within a 100x100 m plot (Sparrow *et al*. 2020b). The percent cover of each vascular plant species was calculated for 629 TERN Ausplots sites with a total of 730 survey visits conducted between 2011—2019, using the point-intercept data (TERN 2020). The 730 surveys included 101 repeat visits carried out with revisit intervals of one to six years (median revisit interval = 4 years). Inter-survey compositional variation (i.e., change in species composition from baseline to revisit surveys) includes noise and trend (see Baruch *et al*. 2022) and its magnitude was used to inform realistic ranges of simulated change parameters (see below). Revisits were subsequently excluded from analysis so that only baseline visits were included in change detection simulations.

**Figure 1.**
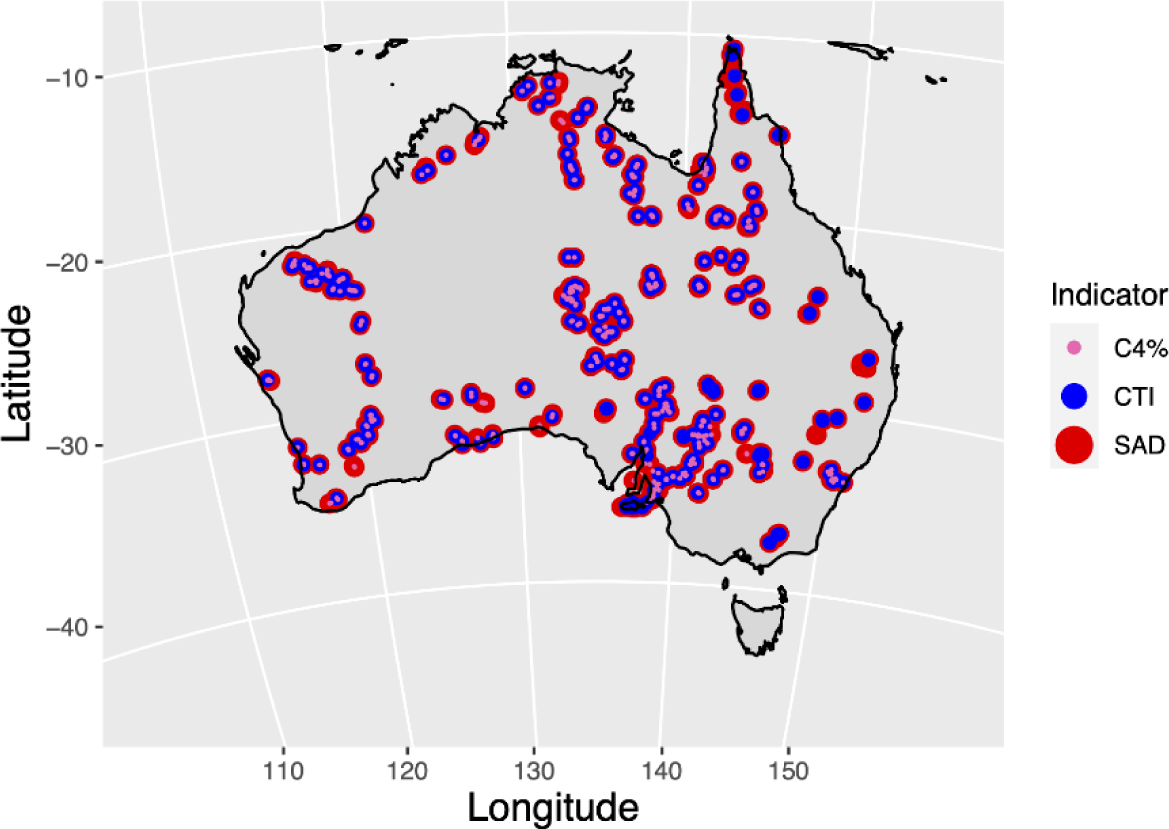
Distribution of TERN Ausplots in Australia that were included in the analysis for the ecological indicators CTI (420 surveys of 382 plots), C_4_% (567 surveys of 527 plots) and SAD (730 surveys of 629 plots) indicators (see legend and Table 1 for indicator definitions).

**Table 1.** Plant community indices based on species composition and relative abundance applied to TERN Ausplots as ecological indicators for change detection. Maximum values for simulated change (and scenario increments) are given.

| Code | Indicator | What does it measure? | Putative drivers | Calculation | Simulated change (increment) | Data reference |
| --- | --- | --- | --- | --- | --- | --- |
| CTI | Community temperature index | Aggregated inferred thermal tolerance | Long-term temperature | Community weighted mean of upper realised (98<sup>th</sup> percentile) thermal limit | 3 (0.1) | Gallagher *et al.* (2019) |
| C<sub>4</sub>% | Photosynthetic pathway | Proportional abundance of C<sub>4</sub> species | Rainfall, temperature, seasonality | Proportion of total plant cover made up of C<sub>4</sub>% photosynthetic pathway species | 0.3 (0.01) | Munroe *et al.* (2021a) |
| SAD | Species abundance distribution | Relative species dominance/evenness | Rainfall, disturbance | Sigma parameter of the lognormal distribution | 0.5 (0.02) | Guerin *et al.* (2017) |

### Indicators

We selected the community temperature index (CTI), the proportional abundance of C_4_ species (C_4_%), and the species abundance distribution (SAD) as indicators of key functional shifts mediated by species composition responses to rainfall, temperature or disturbance. These indices, while based on raw species relative abundance data, are themselves species-free (Table 1).

The CTI is based on empirical realised thermal limits calculated by examining the known range of the species from occurrence data (Gallagher *et al*. 2019). CTI is the abundance-weighted community mean of species temperature preferences (Bowler & Böhning-Gaese 2017), here implemented as upper thermal limit (98^th^ percentile of recorded occurrences with respect to mean annual temperature). While CTI therefore does not encode species physiological tolerances per se, which may be broader (Bush *et al*. 2017), it is nevertheless a useful climatic correlate of expressed species composition, which may represent physiological limits or thermal niche truncation due to competition or other factors. CTI can be compared to the long-term average climate conditions at the sites to assess ‘safety margin’ (CTI minus site mean) and vulnerability (safety margin minus exposure to climate change) to see where change is most likely to occur (Gallagher *et al*. 2019), though interpretation of climatic niches must be made with caution given the importance of demographic and historical factors in determining observed distributions (Fordham *et al*. 2012). In addition to spatial surveys to estimate climate vulnerability across a set of communities, the CTI can also be monitored through repeat visits to detect climatic signals in species composition changes (Kampichler *et al*. 2012). Ideally, directional change in CTI through time across a set of spatially replicated plots will not occur due to ‘noise’ (e.g., sampling error, local disturbance), which should be random with respect to thermal tolerances. However, confounded trends in CTI are still possible where other functional responses correlate with those related to climate, or when an apparent trend in the CTI is driven by a ubiquitous species that has changed its abundance due to some other factor. Such confounding changes, however, become increasingly unlikely when CTI change is detected across larger areas and different species assemblages.

For CTI, climatic niche data were sourced from Gallagher *et al*. (2019) and consisted of 98^th^ percentiles of mean annual temperature across species recorded ranges based on intersecting filtered herbarium records with gridded long-term climate datasets from WorldClim (Hijmans *et al*. 2005). To calculate CTI, we merged species-level climatic niche data with the species composition matrix, which was trimmed to include species represented in the climate niche data, then subset to sites with at least 80% coverage by abundance (Pakeman & Quested 2007; Borgy *et al*. 2017; Guerin *et al*. 2021b). CTI was calculated as the community weighted mean (Hulshof *et al*. 2013) of the maxima (98^th^ percentile) of species niches for mean annual temperature and has units of degrees Celsius.

The C_4_% index is the proportional abundance of C_4_ (cumulative across species) relative to C_3_ and CAM photosynthetic pathways in a vegetation community and reflects photosynthetic efficiency in different environments (Munroe *et al*. 2021a). The C_4_% is therefore expected to be a strong indicator of response to temperature and rainfall redistribution under climate change, as well as an important indicator of functional changes, such as carbon assimilation and water use efficiency (Lavorel & Garnier 2002; Munroe *et al*. 2021a). Photosynthetic pathway was matched to species records based on an existing compilation which targets species in the Ausplots dataset (Munroe *et al*. 2021a; Falster *et al*. 2021). C_4_% is based on Munroe *et al*. (2021a) and calculated as the proportional cover of species assigned as C_4_ photosynthetic pathway out of the total cover of all species, and so ranges from 0–1.

The SAD is based on Whittaker plots (ranked log-abundance plots; Whittaker 1965), which are fitted to a distribution. The fitted shape parameter relates to the degree to which species abundances are uneven and is a putative indicator of community responses to climate and disturbance that reflect ecological dominance (McGill *et al*. 2007; Guerin *et al*. 2017). Using SAD approaches, evenness in tree assemblages globally has been shown to be correlated positively with actual evapotranspiration and negatively with climatic variability (Ulrich *et al*. 2016). SADs can be calculated directly from species abundance data. For the SAD indicator, we fitted a lognormal distribution to a vector of species abundances at each plot and extracted the sigma parameter (standard deviation, in units of log-transformed abundance).

Because of differences in species coverage among the three indicators, a different subset of the matrix was used for each, hence 382, 527, and 629 baseline plots were used for CTI, C_4_% and SAD, respectively. The different plots and species included per indicator place some limitations on comparisons.

### Power analysis

We estimated power to detect changes in the three indicators across a set of spatial clusters of plots as a model for detecting real trends in these indicators occurring at regional level where broad climatic responses are likely to be evident and at which level the plot network is replicated. Spatial clusters were determined for each indicator separately (due to inclusion of different subsets of available plots) by calculating pairwise geodesic distances among the plot locations and then performing *k*-means clustering on those distances, a method which partitions groups to minimise within-group variance with a specified number of groups but no constraint on group size (Hartigan & Wong 1979). The number of groups (30 for CTI, 40 for each of C_4_% and SAD) was selected to produce well-defined geographic clusters given the number of plots available. Note that the inclusion of different plots and the use of different clustering methods would generate different clusters which may affect the results of power analysis for a given location. For example, clusters casting larger geographic nets may capture more plots but also more standing spatial variation (Fig. S6).

We simulated change scenarios for each cluster separately that each consisted of a directional change increment to the indicator, accompanied by a random term representing individual plot-level change and sampling noise that differs by indicator and scenario but not by cluster (Eq. 1; Table 1). For a given plot, the simulated response for each of 1,000 random noise replicates is given by:

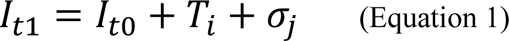

where *I_t1_* is the simulated value of the indicator at time 1, *I_t0_* is the observed empirical value of the indicator at time 0, *T_i_* is the change increment in trend scenario *I* (applied per cluster), and *σ_j_* is a plot-specific random term drawn from a normal distribution with mean zero and standard deviation *j*, which, depending on scenario, ranges from zero to a maximum representing the standard deviation of the indicator among all plots in the baseline dataset.

For each spatial cluster and scenario of change increment, then, there were 1,000 random noise replicates per noise level. At each iteration, change detection in a given cluster was tested by extracting the p-value of a linear mixed-effects model in which plot was the random effect (to account for non-independence between a given plot’s baseline and simulated revisit) and time 0/1 was the fixed effect in predicting the indicator. Because C_4_% is a proportion but not strictly binomial, logit-transformed linear models were applied in that case (Warton & Hui 2011). For a given cluster/change/noise scenario, power was scored as the proportion of the 1,000 replicates in which the change was detected at p < .05 (O’Hare *et al*. 2020). Additionally, we examined variation in the effect size of the directional change (given a noise scenario) by extracting the marginal pseudo R-squared of the linear mixed effects model using the method of Nakagawa *et al*. (2013) as implemented in Bartoń (2020). Effect size was scored as the mean of this statistic over the 1,000 replicates for each scenario.

We visualised and further investigated patterns of power in relation to the change scenario using power curves (O’Hare *et al*. 2021; Figs 2–4) and characteristics of the spatial clusters, such as geographic location, number of plots in the cluster and variance of the indicators in baseline surveys among the plots, as measured through standard deviation (Fig. 5). Power curves were used to extract minimum detectable thresholds of change defined as the smallest magnitude trend for which power was at least 0.8 (Table 2).

**Figure 2.**
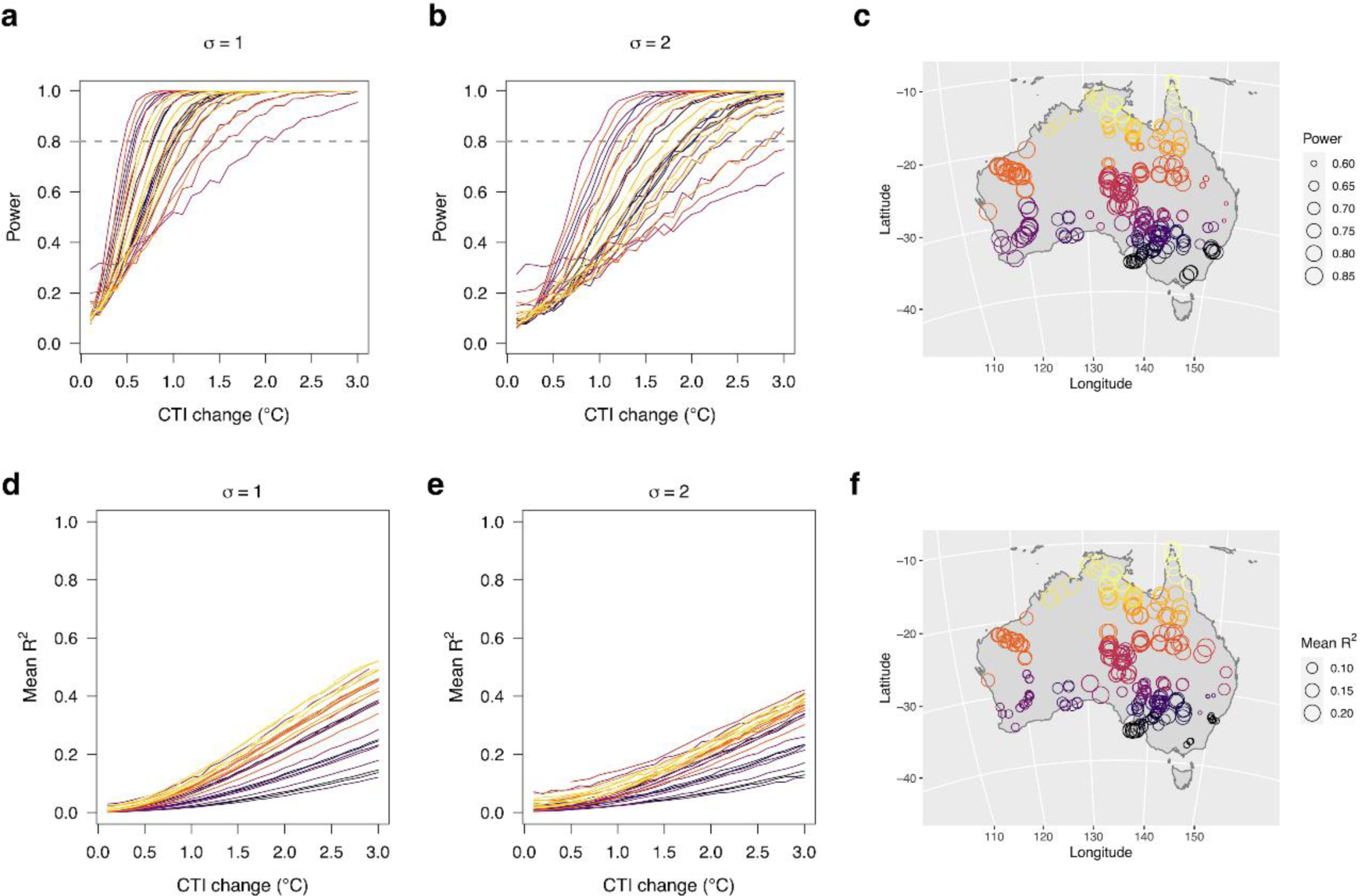
Community Temperature Index (CTI). **a–b)**: Power to detect increases in CTI across 30 spatial clusters of TERN Ausplots (coloured lines corresponding to latitude in maps). Two ‘noise’ scenarios are depicted where σ = standard deviation of a random normal distribution, representing plot-to-plot species composition dynamics due to season of observations, local disturbance, neutral fluctuations or sampling error. Power is calculated as the proportion of 1,000 simulations per scenario in which a linear mixed effects model detected the change at p < .05. A power of 0.8 (dashed lines) is the target minimum. This corresponds to detectable trends of (0.5–) 0.63–2.3 (– 3.0)°C in CTI in format (a–) b–c (–d), where a: minimum when σ = 1; b: σ = 1 & z-score = -1; c: σ = 2 & z-score = 1; d: maximum when σ = 2 (see Table 2), while two clusters did not reach power in b); **c)** spatial plot clusters (circles coloured by mean cluster latitude corresponding to line plots) with point size proportional to mean power. Clusters were assigned using k-means clustering on geographic distances among plot locations, partitioning groups to minimise sums of squared distances to group centres; **d–e)**: Effect size of increases in CTI across clusters with two noise scenarios. Effect size is calculated as the mean marginal pseudo-R^2^ of linear mixed effects models across 1,000 simulations per scenario; **f)**: spatial plot clusters with point size proportional to mean effect size.

**Figure 3.**
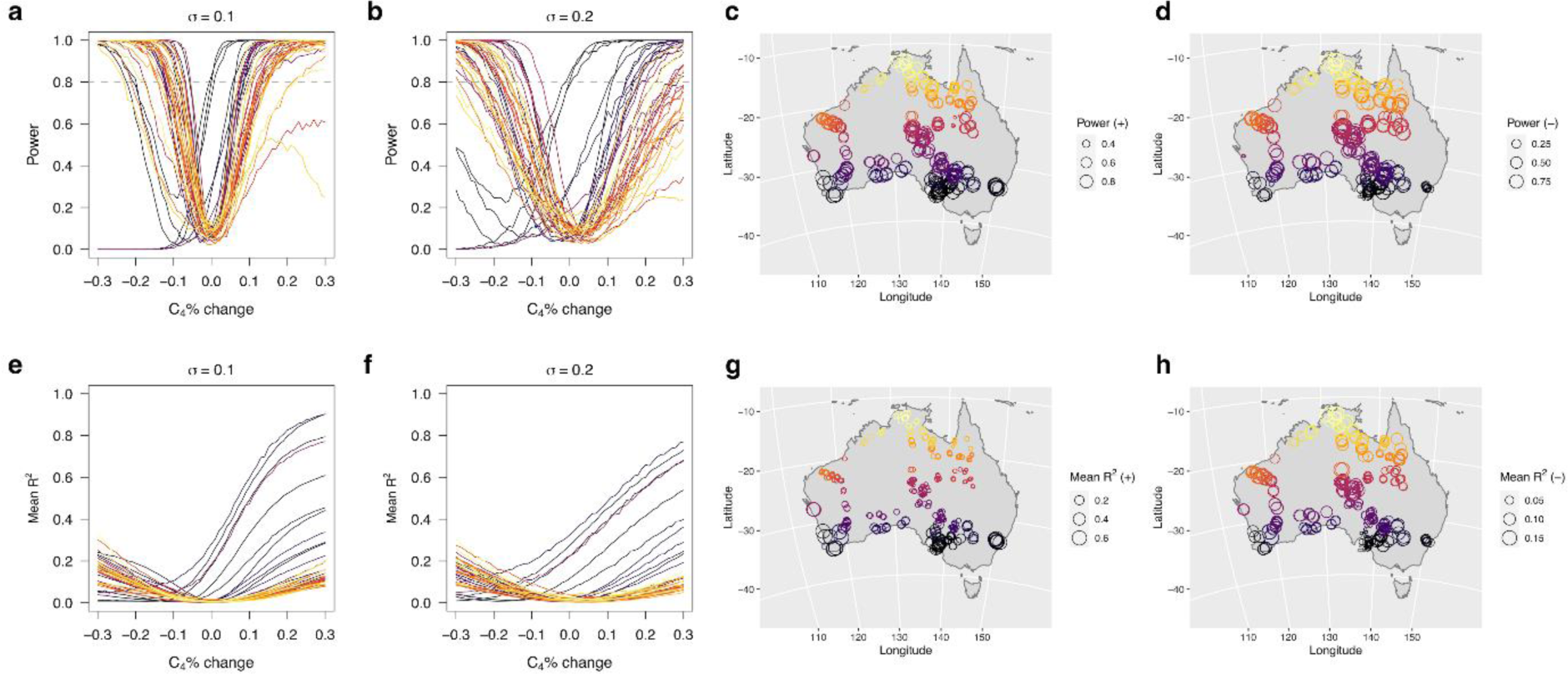
C_4_%. **a–b)**: Power to detect trends in C_4_% across 40 spatial clusters of TERN Ausplots (coloured lines corresponding to latitude in maps). Two ‘noise’ scenarios are depicted where σ = standard deviation of a random normal distribution. Power is calculated as the proportion of 1,000 simulations per scenario in which a linear mixed effects model detected the change at p < .05. A power of 0.8 (dashed lines) is the target minimum. This corresponds to detectable increasing trends of (0.06–) 0.07–0.24 (–0.3) in C_4_% and decreasing trends of (0.01–) 0.06–0.27 (– 0.29), in format (a–) b–c (–d), where a: minimum when σ = 1; b: σ = 1 & z-score = -1; c: σ = 2 & z-score = 1; d: maximum when σ = 2 (see Table 2); **c–d)** spatial plot clusters (circles coloured by mean cluster latitude corresponding to line plots) with point size proportional to mean power for increasing (c) and decreasing (d) trends; **e–f)**: Effect size of trends in C_4_% across clusters with two noise scenarios. Effect size is calculated as the mean marginal pseudo-R^2^ of linear mixed effects models across 1,000 simulations per scenario; **g–h)**: spatial plot clusters with point size proportional to mean effect size for increasing (g) and decreasing (h) trends.

**Figure 4.**
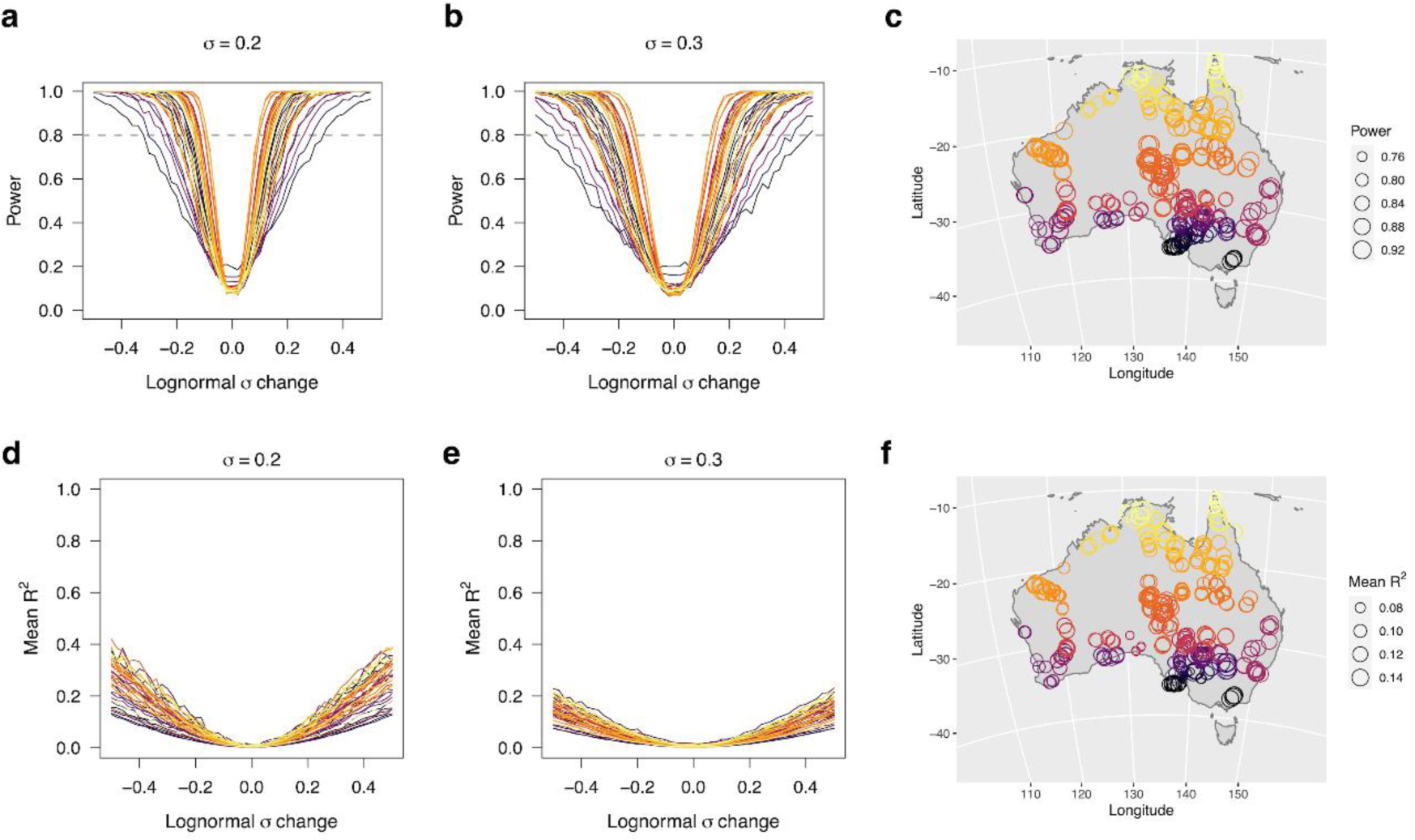
Species Abundance Distribution (SAD). **a–b)**: Power to detect trends in SAD across 40 spatial clusters of TERN Ausplots (coloured lines corresponding to latitude in maps). Two ‘noise’ scenarios are depicted where σ = standard deviation of a random normal distribution. Power is calculated as the proportion of 1,000 simulations per scenario in which a linear mixed effects model detected the change at p < .05. A power of 0.8 (dashed lines) is the target minimum. This corresponds to detectable increasing trends of (0.10–) 0.12–0.34 (–0.5) in SAD and decreasing trends of (0.10–) 0.12–0.33 (–0.48), in format (a–) b–c (–d), where a: minimum when σ = 1; b: σ = 1 & z-score = -1; c: σ = 2 & z-score = 1; d: maximum when σ = 2 (see Table 2); **c)** spatial plot clusters (circles coloured by mean cluster latitude corresponding to line plots) with point size proportional to mean power.; **d–e)**: Effect size of trends in SAD across clusters with two noise scenarios. Effect size is calculated as the mean marginal pseudo-R^2^ of linear mixed effects models across 1,000 simulations per scenario; **f)**: spatial plot clusters with point size proportional to mean effect size.

**Figure 5.**
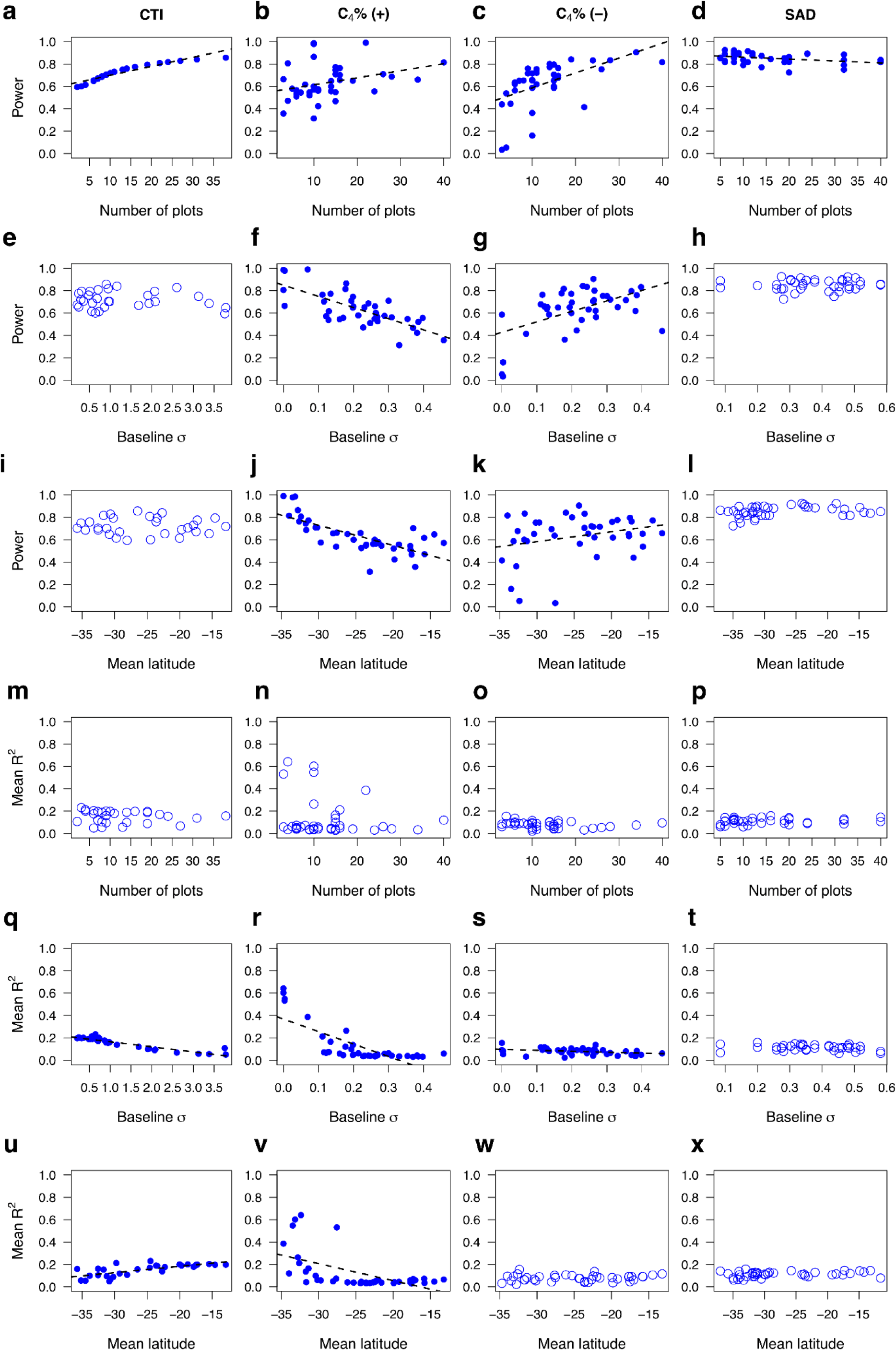
Pairwise associations of mean power and effect size (as ‘mean R^2^’) with number of replicate plots, baseline σ, and latitude at cluster level for the ecological indicators CTI, C_4_% (increasing (+)/decreasing (-) trends) and SAD across the full set of change (trend and noise) scenario simulations explored for TERN Ausplots. Significant linear models indicated with dashed lines.

**Table 2.** Thresholds for detection of change in three ecological indicators across TERN Ausplots, defined as the minimum trend detectable in a spatial cluster of plots with power of at least 0.8. The mean and range of threshold values (as absolute difference) over all clusters are given. Two of the intermediate noise levels that were simulated are displayed per indicator below as examples. See Table 1 for indicator definitions.

| Indicator | Noise SD | Threshold<br>(mean ± SD) | Threshold<br>(range) |
| --- | --- | --- | --- |
| CTI | 1 | 0.98 ± 0.35 | 0.5–2.1 |
| CTI | 2 | 1.8 ± 0.52 | 1.0–3.0 |
| C<sub>4</sub>% (decreasing) | 0.1 | 0.11 ± 0.05 | 0.01–0.24 |
| C<sub>4</sub>% (decreasing) | 0.2 | 0.18 ± 0.09 | 0.01–0.29 |
| C<sub>4</sub>% (increasing) | 0.1 | 0.12 ± 0.05 | 0.06–0.23 |
| C<sub>4</sub>% (increasing) | 0.2 | 0.18 ± 0.06 | 0.01–0.30 |
| SAD (decreasing) | 0.2 | 0.17 ± 0.05 | 0.1–0.32 |
| SAD (decreasing) | 0.3 | 0.25 ± 0.08 | 0.14–0.48 |
| SAD (increasing) | 0.2 | 0.17 ± 0.05 | 0.10–0.34 |
| SAD (increasing) | 0.3 | 0.26 ± 0.08 | 0.14–0.50 |

Analyses were conducted in R versions 3.4 and 3.6 (R Core Team 2018; 2020) using packages ausplotsR, car, geosphere, lme4, MuMIn, parallel, pwr, sads, vegan (Bates *et al*. 2015; Prado *et al*. 2018; Fox & Weisberg 2019; Bartoń 2020; Champely 2020; Oksanen *et al*. 2020; Guerin *et al*. 2021c; Hijmans 2021; Munroe *et al*. 2021b).

## Results

### CTI

Power to detect change in CTI was related to trend magnitude (positive association), noise standard deviation (negative association) and identity of the cluster of spatial plots (Fig. 2A, B). Detectable thresholds of change varied by cluster identity from 0.5–3.0°C for intermediate noise scenarios (Table 2). Mean power across all simulations for a given cluster was positively correlated with the number of plots in the cluster (Fig. 5A) but unrelated to the standard deviation of baseline measurements (Fig. 5E). The relationship of power with spatial replication was curvilinear, attenuating with replication over approximately 15 plots. The spatial arrangement of clusters that had high power reflects baseline sampling intensity, due to the tight relationship with replication, and therefore has no particular geographic pattern (Figs 2C, 5I). Effect size also differed by cluster identity for a given scenario (Fig. 2D, E), and was negatively correlated with the standard deviation of baseline measurements (Fig. 5Q). This translated to a continental pattern of variation in which effect size was lower for more southern plots, which matched a pattern of lower baseline standard deviation in CTI north of 25°S (Figs 2F, 5U).

### C_4_%

Power to detect change in C_4_% was related to trend absolute magnitude (positive association), noise standard deviation (negative association) and identity of spatial plot cluster (Fig. 3A, B). Power also differed between increasing and decreasing trends by cluster identity. Detectable thresholds of change under intermediate noise scenarios varied by cluster identity from 0.01–0.3 (Table 2). For increasing C_4_% trends, mean power had a positive relationship with the number of plots in the cluster (Fig. 5B), a negative relationship with standard deviation of baseline measurements among replicate plots (Fig. 5F), and a negative association with latitude (degrees north), with higher power to detect increases in C_4_% to the south (Fig. 5J). For decreasing trends, mean power increased with the number of spatial replicate plots (Fig. 5C). Mean power also increased with the baseline standard deviation of clusters (Fig. 5G) because southern plots that were already uniformly low in C_4_% at baseline had low power to detect further decreases. Effect size also differed by cluster identity and direction of trend, with some clusters better able to detect increases than decreases and vice versa (Fig. 3E, F). Effect size decreased with increasing standard deviation of baseline measurements (Fig. 5R, S), and was therefore higher along the southern margin of the continent below a latitude of -30N for increasing trends, where plots were uniformly low in C_4_ (Figs 3G, 5V). In the case of decreasing trends, there was no spatial pattern of effect size (Figs 3H, 5W).

### SAD

Power to detect change in SAD was related to trend absolute magnitude (positive association), noise standard deviation (negative association), and identity of cluster (Fig. 4A, B). Detectable thresholds of change with intermediate noise scenarios varied by cluster identity from 0.1–0.5 (Table 2). Mean power across all simulations was consistently high but showed a significant but very slight negative relationship with number of spatial replicate plots (Fig. 5D). Power was not associated with standard deviation of baseline measurements (Fig. 5H), and showed no spatial pattern (Figs 4C, 5I). Effect size increased with absolute trend magnitude and decreased with noise standard deviation (Fig. 4D, E). There was no relationship of effect size with baseline spatial replication, standard deviation, nor was there any spatial pattern (Figs 4F, 5P, T, X).

## Discussion

Monitoring networks are faced with competing priorities. Given finite resources for additional surveys, there is a balance between adding replicates to a sampled ecosystem on one hand (resulting in lower spatial variation among plots in a cluster and improved change detection with high power), and increasing spatial and environmental representation on the other (Guerin *et al*. 2021a; Martín-Forés *et al*. 2021). This trade-off in sampling extent versus intensity captures the interplay between biological survey (i.e., spatial survey capturing ecological diversity) and monitoring (i.e., temporal survey measuring ecological change), which are combined to some extent in national-scale monitoring networks (Barnett *et al*. 2019). Variation among spatial replicate plot samples of a given ecosystem or regional habitat is due partly to the intrinsic patchiness of the ecosystems being surveyed, and partly to sampling strategy and the relative focus on replication versus capture of variation in vegetation types.

The key point is that while either sampling configuration may be valid, depending on the aim, power to detect change at smaller thresholds comes at the expense of wider representation of ecological diversity and vice versa. Such trade-offs also apply to prioritisation of existing sites for revisits (Martín-Forés *et al*. 2021). By ranking clusters of plots by their contribution to species representation, it is possible to reduce the number of clusters revisited and focus resources on ensuring sufficient replication and more frequent revisits to a selected subset for enhanced change detection.

### Maximising change detection – general recommendations for plot networks

#### 1. Select the unit of change detection

Decisions on the unit of analysis for change detection depend on the target spatial scale and level of biological organisation. The arrangement of plots established in the field, or post hoc grouping for the purposes of analysis, can be configured differently to target either spatial regions or particular habitat types, depending on aims. Purely spatial arrangements can confound local differences in vegetation type, leading to excessive sample variance, and vice versa; grouping by vegetation type alone confounds spatial variation, while post hoc grouping by both space and vegetation type results in smaller numbers of replicates and reduced power. Where spatial clusters of plots are too variable, it may be worthwhile splitting them into habitat-specific subunits which are then supplemented with additional replication (Oliva *et al*. 2020). It is important for change detection, however, that groups of replicate plots (at some level of spatial or biological organisation) are considered the unit of measurement, because change at the individual plot level cannot be extrapolated, while replication along bioclimatic transects is an alternative sampling regime (Gillison & Brewer 1985). We recommend that additional plots boost replicate sampling of similar habitats in a given area to maximise change detection, once the overall representation and inclusivity goals of the network are met (Guerin *et al*. 2020). Here, we applied objective spatial clusters as the units of change detection for regions in Australia because TERN Ausplots has no formal replication of target ecological communities in its sampling strategy, while there is spatial replication by default because plots are established in regional surveys.

In addition to selecting plot samples to form the units of change detection, change needs to be tracked using appropriate ecological indicators (or in the case of TERN Ausplots, measured vegetation parameters). Whereas metrics such as beta diversity rely on pairwise comparisons of shared species among sets of sites, species-free indicators like those employed here can be used to compare community samples more widely through space and time, because samples that share no species do not default to dissimilarities of one, for example. Compared to multivariate approaches to analysing species composition and function (e.g., RLQ, fourth corner, constrained and unconstrained ordination, generalised linear mixed models and so forth; Pakeman 2011), species-free indicators simplify interpretation because they are univariate (i.e., a single value represents each community sample), while representing key functional aspects of the systems (i.e., dominance, thermal tolerance and photosynthetic capacity for the indicators applied in this study).

#### 2. Sufficient replication

Our results highlight the importance of adequate, targeted spatial replication to achieving change detection power in a monitoring network, where the aim is to detect trends at the regional level. Directional change may be modest and masked by plot-to-plot ‘noise’ caused by factors such as local disturbance, non-deterministic (neutral) fluctuations in relative species abundance, or sampling error. The relationship between replication and power suggests additional plots are more effectively placed in clusters with the fewest existing plots, while there can be a law of diminishing returns at higher sampling intensities. Replication to a minimum of 10–15 plots and minimising variance are strategies likely to improve change detection, notwithstanding that power to detect changes in SAD appeared robust to levels of replication. Additional plots may be warranted to increase representation of vegetation types or disturbance levels within a region, or to detect directional change with a small magnitude. Alternatively, plot clusters targeted for revisits could be further vetted based on their ability to be both representative and well-placed for change detection in terms of sufficient replication and modest baseline variance.

#### 3. Maintain species coverage of indicators

Species composition is a key property of the vegetation that is commonly targeted in survey and monitoring protocols (Bellingham *et al*. 2020). While species composition is a complex and labile response with many drivers (Baruch *et al*. 2022), it can be used effectively to underly the calculation of indicators that relate to specific drivers. For example, C_4_% is based on species relative abundance yet is both a response and effect trait in terms of ecosystem functioning (Munroe *et al*. 2021a), while CTI may reveal underlying species composition trends related to climate warming (Bowler & Böhning-Gaese 2017). Application of functional indicators can be limited by incomplete species coverage. Further gap-filling and targeted trait assays on an on-going basis will ensure high coverage and broad utility of trait-based indicators. The indicator SAD uses intrinsic species abundances to create a univariate metric. Although SADs produced less clear patterns here in terms of the relationship of sampling to power, its potential links to disturbance regimes and environmental gradients make further evaluation worthwhile (McGill *et al*. 2007; Guerin *et al*. 2017). There is still significant debate about ecological interpretation (McGill *et al*. 2007) and quantitative trends that reach biological significance in terms of surveillance monitoring are yet to be determined.

#### 4. Minimise noise

Our simulations explored scenarios of random non-trend noise, drawn from distributions with increasing standard deviation. These simulations showed that noise levels are a key factor in determining minimum change thresholds for detectable trends (Table 2). The magnitude of simulated detectable differences was comparable to observed temporal changes, suggesting that our framework for change detection applied retrospectively to the TERN Ausplots network is capable of tracking biologically relevant change when these ecological indicators are applied to the complex species composition data, with the caveat that the ecological response would, in order to correspond to simulated power levels, have to be occurring consistently across all the ecological communities sampled within a regional cluster of plots. Indeed, it is useful to detect trends in the order of 1°C in CTI and 0.1 in C_4_%, as such changes are linked to significant shifts in species composition (Hoffmann *et al*. 2019; Guerin *et al*. 2019) and the water use efficiency of photosynthesis at community level (Munroe *et al*. 2022a; Munroe *et al*. 2022b), respectively. For those clusters with higher detection thresholds or higher potential noise levels, change detection should be shored up with additional replication (see 2. above). Detecting regional trends in the raw species composition data rather than in functional indicators may be less achievable given the variation in vegetation types captured in the TERN Ausplots baseline surveys combined with labile species responses that may be site specific (Baruch *et al*. 2018; Baruch *et al*. 2022).

In practice, one could include additional sampling, analytical controls, or covariates to constrain tests for regional trends with respect to season, local disturbance (e.g., fire, flood, or grazing) or other factors. For example, field plots should be representative, and located to avoid highly modified habitats and on-going local disturbance and edge effects associated with infrastructure such as roads, unless such disturbance is the object of study. Precise measures of species presence and abundance in the field also increase power by reducing ‘noise’ due to sampling error such as species non-detection (Clarke *et al*. 2012), misidentification or unstable estimates of abundance. Variance among baseline spatial plot replicates will always be present due to the patchiness of vegetation composition. Stratification and targeted replication can keep variance due to sampling strategy to a minimum.

### Factors determining power and effect size

Aspects of the power–sampling relationship are determined purely by statistical principles. For example, power can be calculated precisely if sample size, effect size and significance criterion are known (Cohen 1988; Champely 2020). Indeed, when applied to real-world data from a continental plot network, community temperature index (CTI) displayed neat relationships to sampling. Predictable changes in power were evident with the addition of plot replicates, as were linear increases in observed effect size with decreasing baseline sample variance (Fig. 5A, Q). Specifically, increasing replication up to approximately 15 plots substantially increased power.

Nevertheless, power to detect change in the three ecological indicators we tested had some idiosyncratic patterns with respect to sampling configuration (Fig. 5). Some of the geographic differences in power relate to underlying properties of the ecosystems themselves. For example, there is a south–north transition in the Australian vegetation from a predominance of C_3_ to C_4_ in the herbaceous stratum, centred around a latitude of approximately 30°S (Munroe *et al*. 2022a). For this reason, it should be easier to detect increases in C_4_% at more southern latitudes (Fig. 5I), though transitions may be more likely to occur at mid-latitudes. Effect size for CTI was greater north of the tropic of Capricorn, where variance in CTI among spatial replicate plots was lower, meaning smaller trends should be detectable in the north of Australia. This is especially relevant because plant communities in tropical regions of Australia, and globally, may be the most vulnerable to temperature increases due to constituent species occurring close to their realised upper thermal limits (Krishnaswamy *et al*. 2014; Gallagher *et al*. 2019).

Power to detect changes in species dominance, as measured through the species abundance distribution (SAD), while closely tied to trend magnitude, did not have a strong pattern with respect to sample size or variation among baseline observations, and unlike CTI and C_4_%, had no overall spatial pattern. Instead, power was consistently high, even when replication was low (Fig. 5D). Because of the high power to detect shifts in SAD, we can only make one conclusion regarding appropriate sampling configurations, namely that change in SAD is highly detectable and robust to sampling design. SAD is therefore a useful candidate indicator, especially since it can be calculated directly from species relative abundance data yet can be compared in the absence of overlapping species composition.

Overall, we conclude that change detection capacity for CTI and C_4_% is partly biome-specific, notwithstanding sampling strategies, due to ecosystem differences linked to climate. The differences in power among the three indicators suggest patterns for different responses or indicators not tested here may also need to be explored individually. Based on our results, power to detect shifts was lower on the whole for C_4_% than for CTI (with sufficient replication) and SAD. Effect size was predictably high for CTI when baseline sampling variance was minimised. For this reason, CTI appears suitable for monitoring response to climate, while SAD is suitable for monitoring disturbance and community structure.

### Conclusion and recommendations

Power to detect change in three ecological community indicators linked to climate, disturbance and ecosystem functioning, CTI, C_4_% and SAD, revealed differences relating to both sampling configuration and climate type (temperate, desert and tropical) when applied to TERN Ausplots across Australia. The results suggest that change detection needs to consider not only statistical principles but also regional differences in sensitivity and degree of local spatial variation in a desired indicator. Despite these idiosyncrasies, some themes emerge from our analysis:

1. Regional plot clusters for surveillance (status and trends) monitoring should contain a minimum of 10-15 replicates to ensure adequate power to detect change in a range of ecological indicators.
2. Sampling variance should be minimised by stratifying additional replication to target ecosystems, avoiding incidental, localised disturbance, and timing surveys to avoid intra-annual seasonal variation where possible. Indeed, multiple revisits on a regular schedule are needed to distinguish true long-term trends from interannual climatic variation and ecological drift.
3. Trends are easier to detect when sampling error is low. This can be achieved by maintaining high standards of species vouchering, identification and curation and robust protocols for measuring abundance or cover.
4. Interpretation of change will benefit from further research to empirically link indicators and their trends to climate and disturbance regimes in different structural vegetation types.

## Acknowledgements

We thank TERN supported by the Australian Government via the National Collaborative Research Infrastructure Strategy, NCRIS.

## Statements and Declarations

### Data availability

This study used existing data. TERN Ausplots data are freely available via the ausplotsR R package (Guerin et al. 2021c; Munroe et al. 2021b). Data for CTI were sourced from Gallagher et al. (2019). Photosynthetic pathway data were sourced from Munroe et al. (2021a). Data and code used to run simulations and power analysis are available at https://doi.org/10.25909/22912445.v1 (Guerin et al. 2023).

## Supplementary Information

The online version contains supplementary material:

*Figure S6*

