## Supplementary Figure S6 for "Detecting ecosystem trends in response to climate and disturbance across continental plot networks: a power analysis"

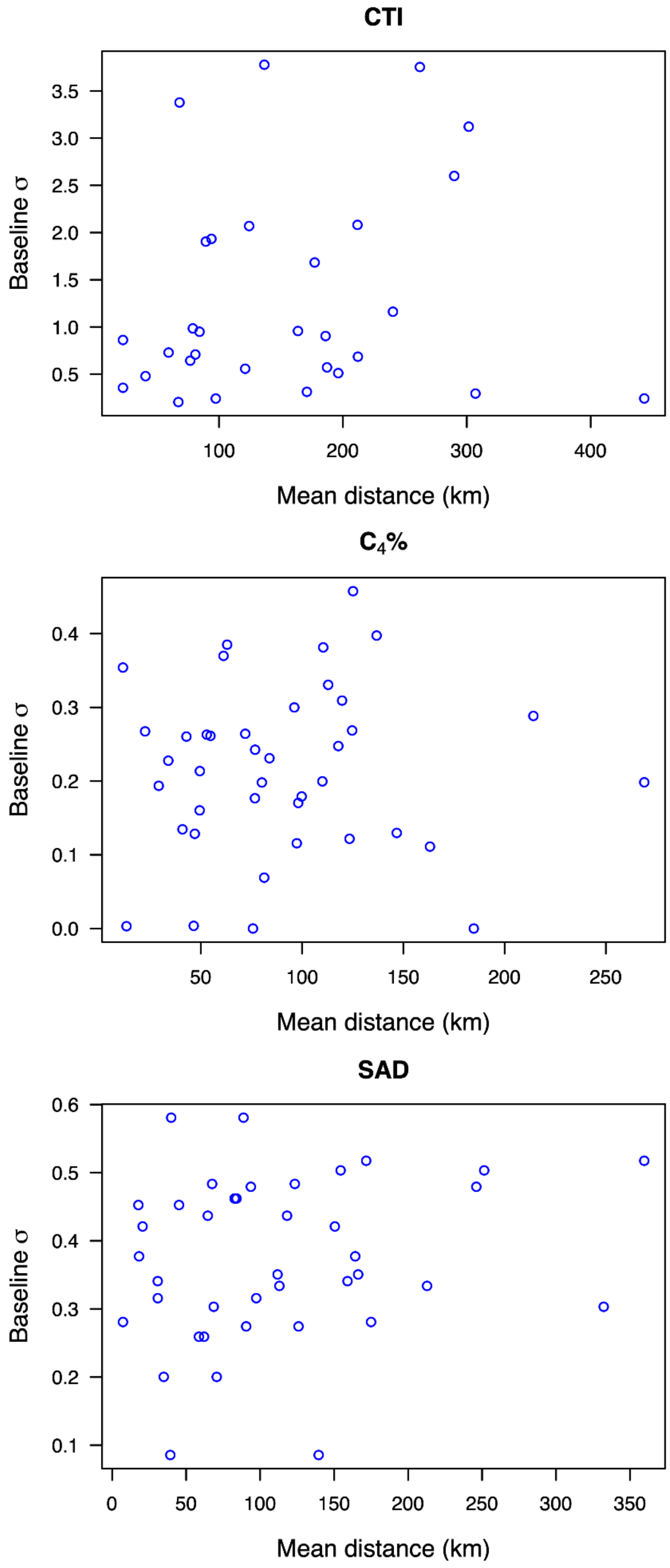


**Figure S6.** Scatterplots showing the relationship between baseline σ (y axis: the standard deviation, representing variation among baseline plots) and the mean geographic distance between plots in a cluster (x axis) for the ecological indicators CTI (Community Temperature Index), $C_4$% and SAD (Species Abundance Distribution) across the TERN Ausplots ecosystem monitoring network. Circles represent individual spatial clusters of between 2 and 40 plots (mean 14 plots). See Table 1 in main text for indicator details. Linear correlations were non-significant in all cases.
